# Finite Element Modeling of Residual Hearing after Cochlear Implant Surgery in Chinchillas

**DOI:** 10.1101/2023.02.15.528713

**Authors:** Nicholas Castle, Junfeng Liang, Matthew Smith, Brett Petersen, Cayman Matson, Tara Eldridge, Ke Zhang, Chung-Hao Lee, Yingtao Liu, Chenkai Dai

## Abstract

Cochlear implant (CI) surgery is one of the most utilized treatments for severe hearing loss. However, the effects of a successful scala tympani insertion on the mechanics of hearing are not yet fully understood. This paper presents a finite element (FE) model of the chinchilla inner ear for studying the interrelationship between the mechanical function and the insertion angle of a CI electrode. This FE model includes a three-chambered cochlea and full vestibular system, accomplished using μ-MRI and μ-CT scanning technology. This model’s first application found minimal loss of residual hearing due to insertion angle after CI surgery and indicates that it is a reliable and helpful tool for future application in CI design, surgical planning, and stimuli setup.

## 1. Introduction

### 1.1. Cochlear electrodes importance and trauma

More than 5% of the world’s population suffers from disabling hearing loss, amounting to over 430 million people [1]. In cases where hearing aids are no longer useful or sufficient, cochlear implant (CI) surgery is the standard procedure for the treatment of severe hearing loss. Modern CI surgery often significantly improves patients’ health-associated quality of life [2,3,4,5,6]. However, it is also known that CI surgery can cause varying levels of trauma or cochlear obstruction that affect the residual mechanical function of the inner ear [7]. The magnitude of this disruption could be partially dependent on the insertion depth of the implant [8,5].

### 1.2. Effect of CI surgery on residual hearing

Most sources report that the magnitude of CI surgery’s effect on residual hearing is largely dependent upon whether cochlear trauma takes place during CI surgery. Trauma is usually attributed to the dislocation of the CI electrode from the scala media or vestibuli [9]. Dislocation can arise due to a variety of factors, thus necessitating the correct choice of an electrode, the surgical technique, and the insertion angle [10]. An important consideration is the morphology of the patient’s cochlea, as shorter and smaller cochleae tend to have higher rates of intracochlear dislocation when fully inserted [11]. In most modern cases, CI electrodes are correctly placed in the scala tympani with minimal trauma. However, a systematic understanding of the effects of typical CI electrode placement on the finer sensitivity of the basilar membrane could be an important step toward further improvement of CI electrode design.

### 1.3. Effect of Insertion angle on CI effectiveness, residual hearing

Insertion angle is a major contributor to CI effectiveness [7]. When longer electrode models are selected, typically with an angle of insertion greater than 540 degrees, lower frequencies become more perceptible to patients. Music becomes more enjoyable, and the quality of life increases compared to those patients with shorter CI electrodes [12]. In cases where CI surgery results in minimal-to-no trauma and the patient is healthy with few underlying conditions, very few side effects have been reported other than postoperative vertigo and nausea [13].

### 1.4. Prior FE models

Over the past decade, substantial research progress has been made to advance inner ear computational modeling. Specifically, finite element (FE) modeling allows the intricacies of the inner ear’s mechanics to be reduced to simpler phenomena that can be verified with clinical results. A previously developed finite element model focused on residual hearing found that cochlear implants most dramatically affect the residual hearing at extreme frequencies of human hearing [14]. However, the previous models did not examine the effect of varying cochlear electrode insertion angles between patients. These unexplored results could provide important metrics for clinical use. Therefore, a comprehensive finite element model capable of simulating hearing function with a variety of insertion angles is an essential step in the improvement of cochlear implant design and surgery.

### 1.5. Chinchilla as an animal model

The chinchilla is used as the animal model in this study. The chinchilla is commonly used as an analog for the human in hearing and balance studies due to its similar number of turns in the cochlea, structure of semicircular canals, singular primary crista, and hearing range [15,16,17]. Chinchillas are commonly used as an analog in FE analysis due to their large bulla and easy availability which allows for very fine resolution of models if scanned with a μ-MRI or μ-CT machine. For the preliminary design of electrode and efficacy measurement, the chinchilla is a good animal model on account of its similarity to human structure, hearing frequency range, low cost, and larger supply source (compared with primates). Furthermore, a chinchilla computational model will be useful to conduct virtual experiments and eventually reduce the extensive use of chinchillas.

### 1.6. Focus of the study

This study focuses on the effect of cochlear electrode insertion depth on the residual mechanical function of the cochlea in a chinchilla computational model. The unimplanted model is demonstrated here first and compared to expected response curves to demonstrate its initial validity. The analysis will then focus on the effects of cochlear implantation on residual hearing.

## 2. Materials and Methods

### 2.1. Data Source and Segmentation

Model geometry was generated through 3D reconstruction of for a single, adult chinchilla. CT scans were acquired at 12 μm voxel size and μ-MRI scans at 30 μm voxel size. Achieving this voxel size with adequate reduction of feedback for the μ-MRI required 26 hours of acquisition in an 11.7 Tesla magnet. The μ-MRI and μ-CT scans of the chinchilla bulla were segmented using a program called 3D Slicer [18]. Images were separated into segments representing lymphatic fluid, bone, and nervous tissue. Sample segmented μ-MRI and μ-CT images are seen in Figures 1 and 2.

**Figure 1.**
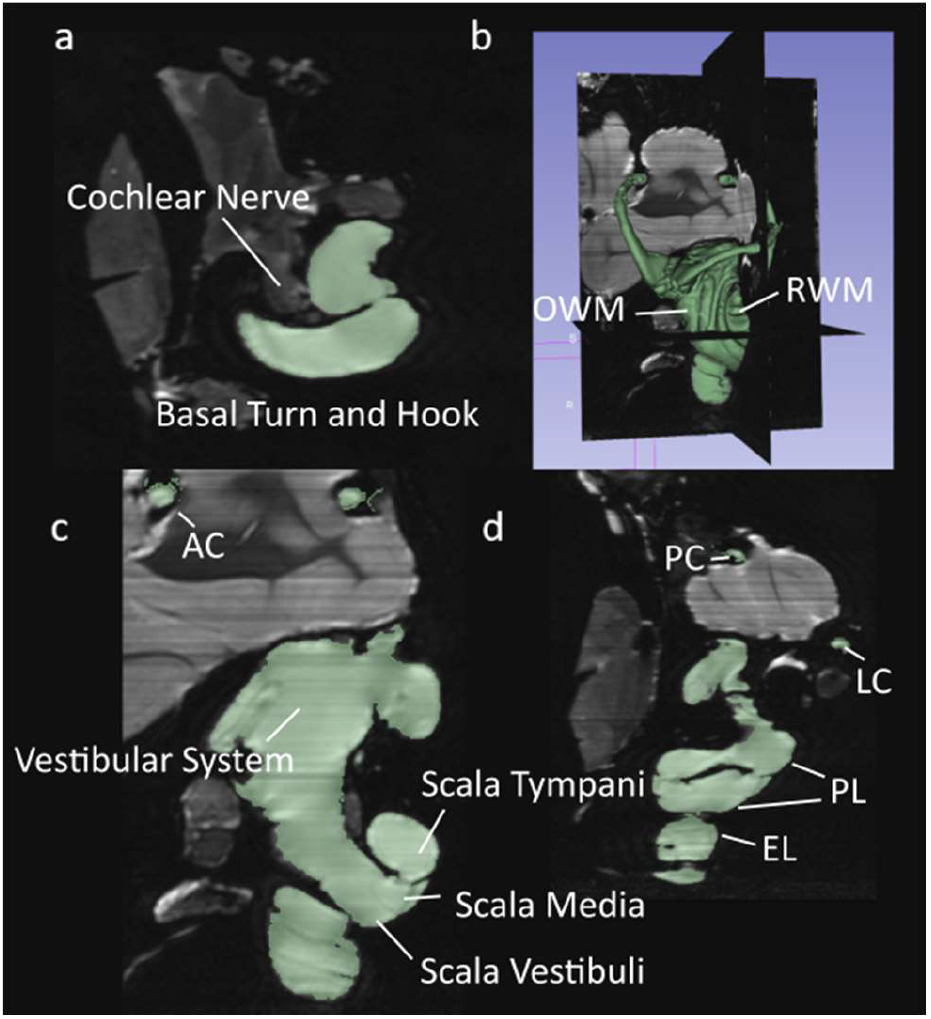
Segmented μMRI scans of the chinchilla subject with key structures labeled. The lymphatic fluid of the inner ear is shown in green. (**a**) Transverse plane (**b**) 3D view of the entire segmentation (**c**) Saggital plane (**d**) Coronal plane. See Table 1 for symbol definitions.

**Figure 2.**
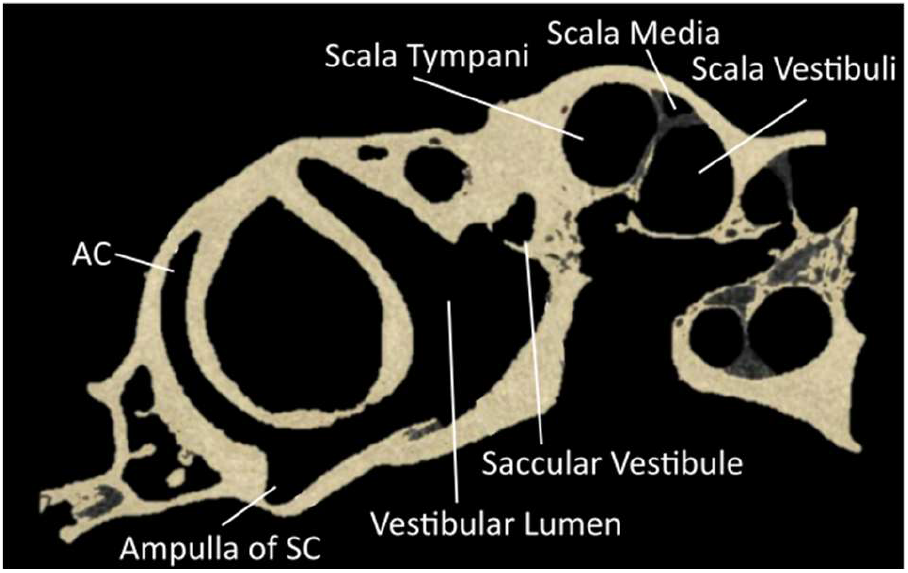
Segmented μCT scan in the sagittal plane of the chinchilla subject with key structures labeled. See Table 1 for symbol definitions.

### 2.2. Geometry

Geometry obtained from 3D Slicer was imported into MeshMixer for the smoothing of surfaces. Numerous small bumps and cavities, due either to bone porosity or imaging artifacts, were removed from the boundary of the bony labyrinth. Holes and gaps in the semicircular canals and cochlea were repaired manually to ensure a continuous volume. Curvatures of repaired sections were matched to those of surrounding surfaces using tools in the MeshMixer program. For this study, all geometry outside the otic capsule was excluded.

The membranous labyrinth of the semicircular canals was modeled in MeshMixer by creating a copy of the bony labyrinth shrunk by a fraction to create two volumes, one enclosed within the other. The utricle’s shape was modified to maintain proper connectivity with the semicircular canals and the ampullae. The utricle was scaled to accommodate a macula consistent with descriptions in literature [19,20]. The saccule was modeled by cross-referencing measurements obtained for humans with data obtained on the saccular macula in the chinchilla. [21,19,22]. The membranous labyrinth in the semicircular canals was scaled to ensure a realistic ratio of endolymphatic fluid to perilymphatic fluid by volume. Cupula structures follow the diaphragmatic model and span the entire width and height of the ampullas. The diaphragmatic model is commonly used in the modeling of vestibular mechanics and yields results that closely mirror reality [23,24,25,26]. The reuniting duct was modeled according to measurements found in literature [27,28,29]. The coordinate system for all figures in this described in Figure 3. An annotated model of the completed vestibular system is illustrated in Figure 4.

**Figure 3.**
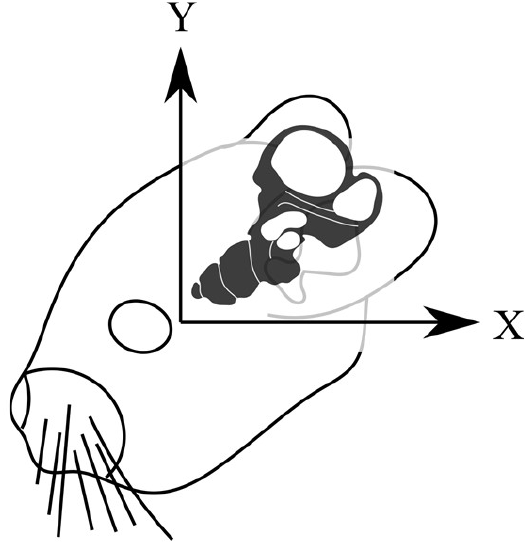
The coordinate system used in all imaging for this paper. The x and z axes are held in the sagittal plane as if viewed from the subjects left side.

**Figure 4.**
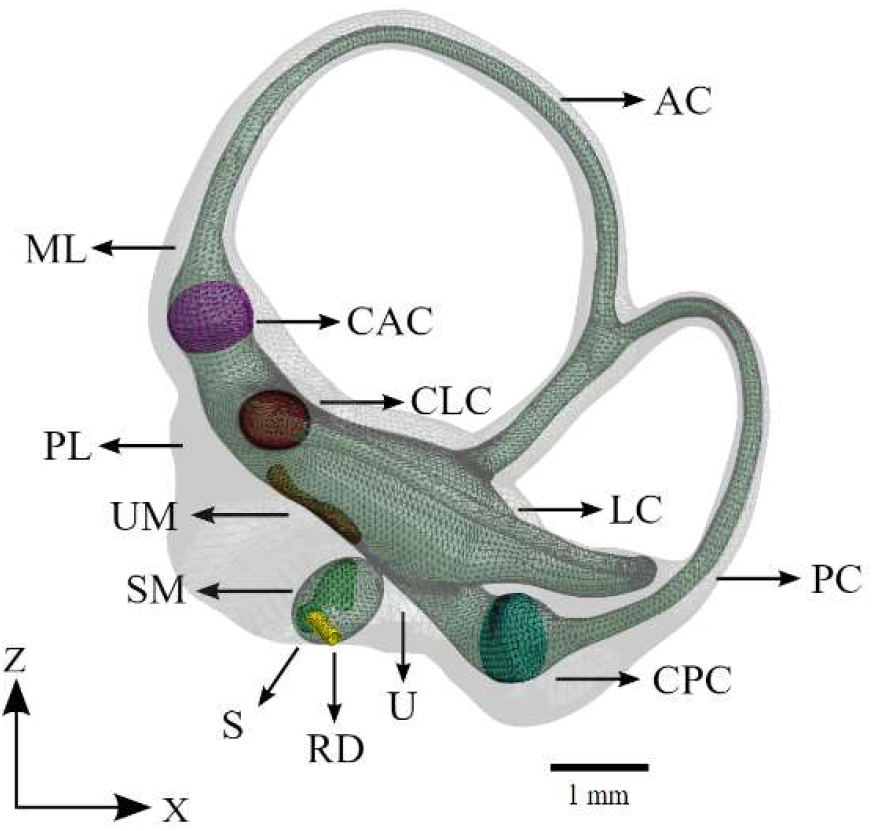
The vestibular system of the computational model. The saccule, utricle, and semicircular canals appear as a continuous volume of lymphatic fluid (green). The sensory organs of the vestibular system are also shown. See Table 1 for symbol definitions.

The cochlea was modeled primarily based on information obtained from μ-MRI imaging. Characteristic ridges on the surface of the bony labyrinth were used to determine the attachment points of the Reissner’s and basilar membranes. The osseous spiral lamina was also clearly defined and marked the inner attachment of both the Reissner’s and basilar membranes. These curves were connected by planes forming a wedge whose superior face represents the Reissner’s membrane and whose inferior face represents the basilar membrane. This shape was compared with that seen in literature and was confirmed to have the correct structure [30]. The completed model of the cochlea is displayed in Figure 5.

**Figure 5.**
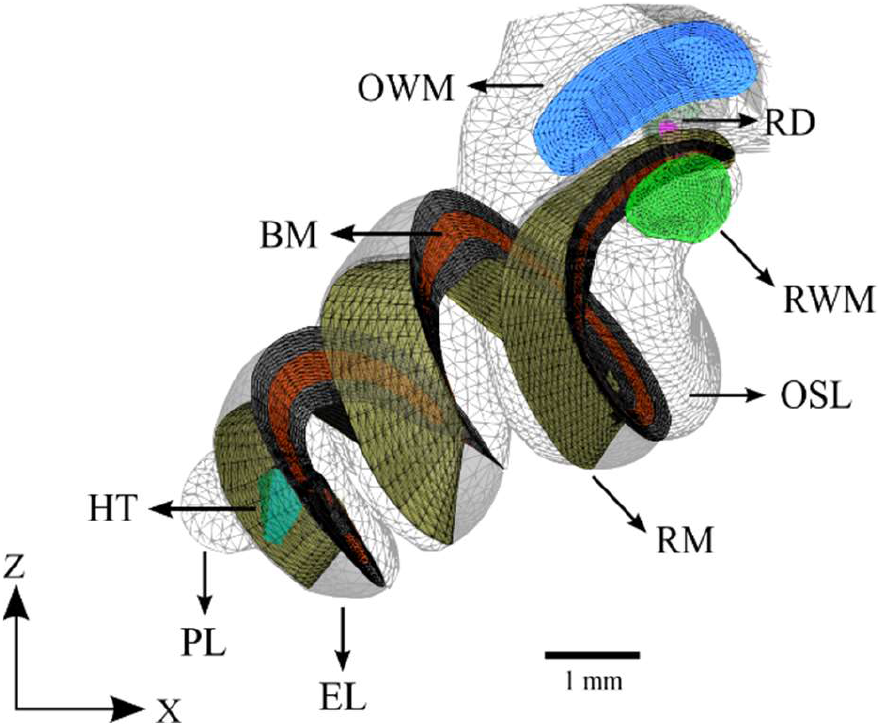
The cochlea of the computational model without cochlear implant. The design of the basilar membrane (red) is apparent as a ribbon with varying width and thickness, attached on both sides to bony supports (grey). The distal side of the basilar membrane is visible as the attachment point for the end of the reissner’s membrane. See Table 1 for symbol definitions.

The cochlea was scaled until the basilar membrane was the average length in chinchillas of 18.3 mm along its midline [31]. The thickness of the basilar membrane was varied from 16.5 μm at the base to 5 μm at the tip according to the values given for the pars pectinate in Cochlear Anatomy and Central Auditory Pathways [32].

Dimensions of a MED-EL FLEXSOFT electrode array were scaled to create an analogous implant which was placed in accordance with an ideal round window insertion in the scala tympani of the cochlea. The cochlear electrode extends almost the full length of the scala tympani with a maximum insertion angle of 900 degrees. This implant was split into 180-degree sections to allow for analysis with varying angles of insertion. Cross sections at the proximal and terminal ends of the cochlear electrode are shown in Figure 6.

**Figure 6.**
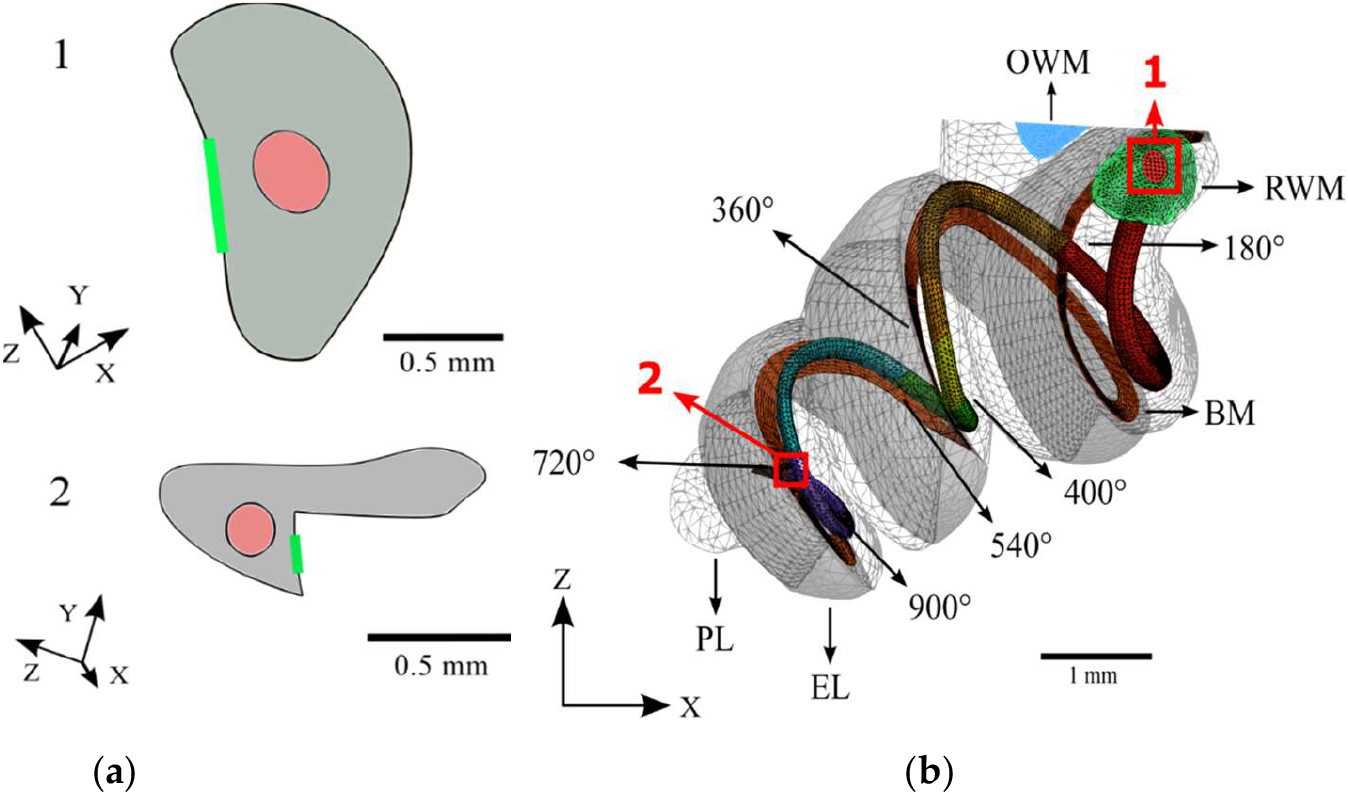
The full meshed model of the cochlea is presented with the length of the cochlear implant electrode inserted. (**a**) Cross sections of the base (1) and apex (2) ends of the cochlear implant electrode; (**b**) The path of the cochlear implant electrode through the scala tympani of the meshed model.

### 2.3. Meshing

Only the components relevant to the mechanical model were meshed. This includes the oval window membrane, round window membrane, cupulas, vestibular maculae, basilar membrane, Reissner’s membrane, utricle, saccule, semicircular canals, cochlea, and cochlear implant electrode array. The mechanical model was meshed with a total of 414,629 tetrahedral elements and 90,696 nodes using the software HyperMesh. Mesh size convergence analysis was not conducted due to the already very fine average element size of about 0.2 mm. This size is sufficient when considering the larger element sizes utilized by previous models [33,34]. However, applying mesh size convergence analysis to future iterations of the model may reduce the processing power and time necessary for simulation [35]. Tissues and the electrode array were modeled using the Ansys Solid185 element type while the fluids were modeled using the Ansys Fluid40 element type. A fine-ruled mesh composed of 8,738 elements was chosen for the basilar membrane. All components were then assigned their respective thicknesses, given proper connectivity, and meshed with tetrahedral elements using HyperMesh.

### 2.4. Material Properties

Due to the relative scarcity of published data on the material properties of chinchilla inner ear soft tissues, material properties measured in humans have been substituted as needed. All material properties are located in Table 2. Mechanical properties of the RWM were gathered from Zhang et al. [36] and Gan et al. [37]. The RM and BM in this model have varied Young’s moduli and damping factors along their lengths and 0.4 as their Poisson’s ratios [38]. The material properties of the endolymph and perilymph were assumed to be identical due to their similar composition. Exponential equations describing these quantities were selected to ensure the model’s results most closely resembled the experimental results that they are compared to as a baseline. These equations were determined through repeat simulation using different plausible functions dependent on position along the cochlea.

**Table 1.**
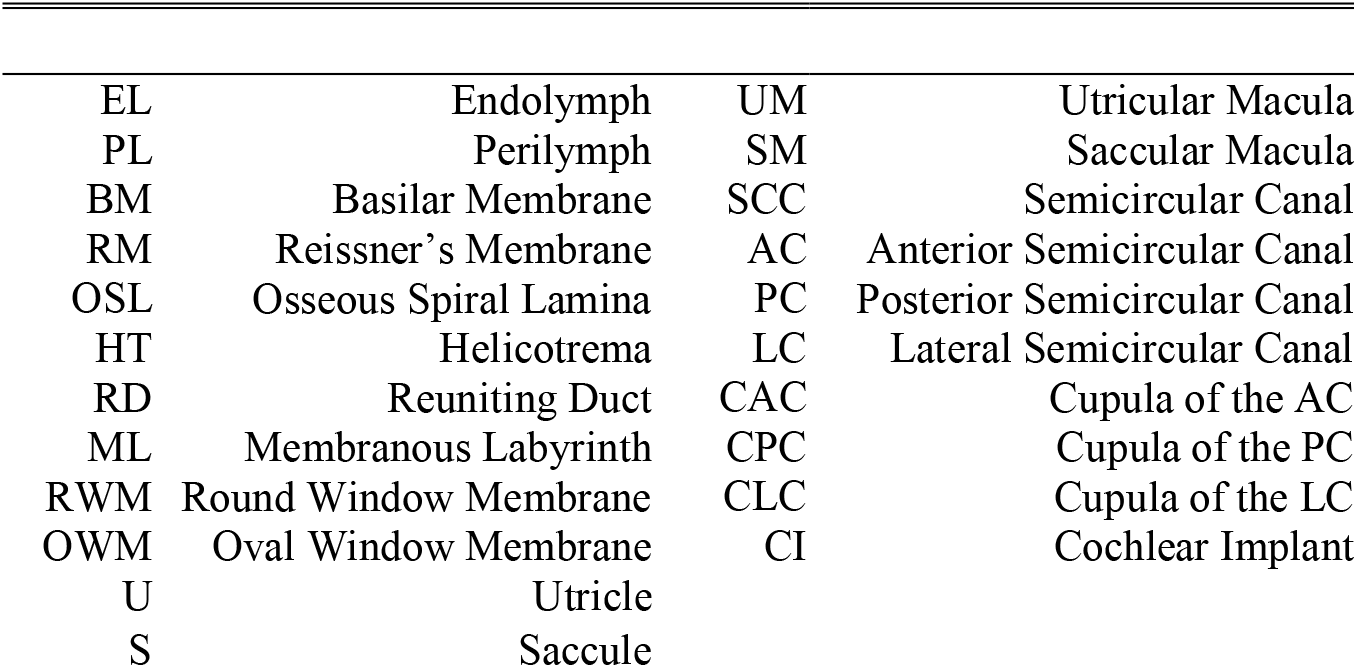
List of Abbreviations

|  |  |  |  |
| --- | --- | --- | --- |
| EL | Endolymph | UM | Utricular Macula |
| PL | Perilymph | SM | Saccular Macula |
| BM | Basilar Membrane | SCC | Semicircular Canal |
| RM | Reissner's Membrane | AC | Anterior Semicircular Canal |
| OSL | Osseous Spiral Lamina | PC | Posterior Semicircular Canal |
| HT | Helicotrema | LC | Lateral Semicircular Canal |
| RD | Reuniting Duct | CAC | Cupula of the AC |
| ML | Membranous Labyrinth | CPC | Cupula of the PC |
| RWM | Round Window Membrane | CLC | Cupula of the LC |
| OWM | Oval Window Membrane | CI | Cochlear Implant |
| U | Utricle |  |  |
| S | Saccule |  |  |

**Table 2.**
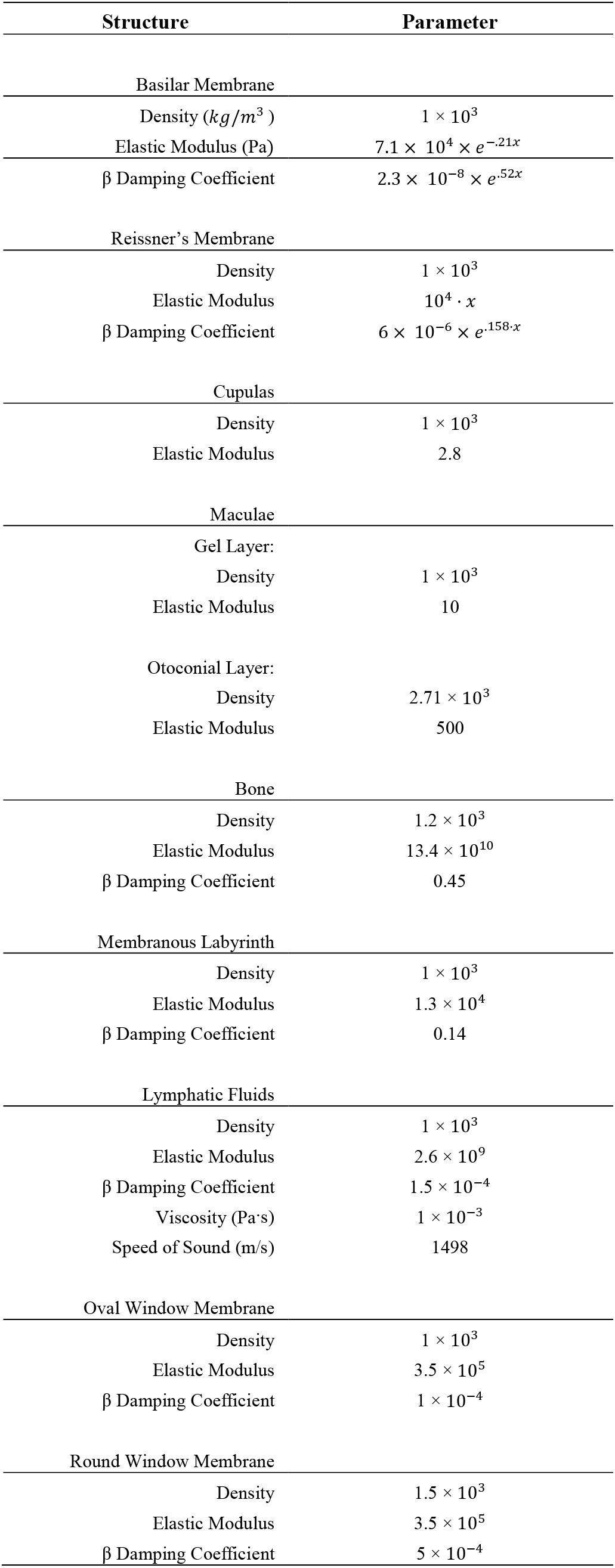
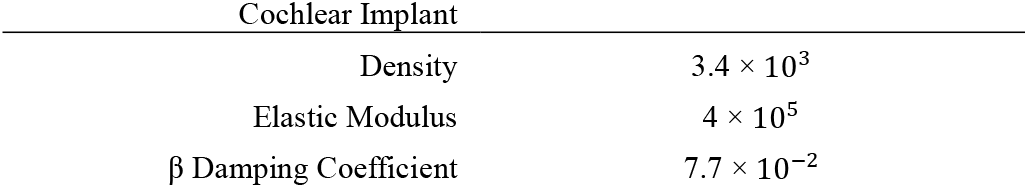
Mechanical Properties of the Model

The material properties of the cochlear implant electrode array were based on the Nucleus Straight electrode array as the material properties of the MED-EL models are not published. A Young’s modulus of 0.4 MPa and density of 3400 kg/m3 were used as has been previously done by Lim et al. in their finite element model of residual hearing after cochlear implantation [14]. The damping factor of cochlear implant electrode arrays has not been published and was thus assumed to be that of the carrier material, silicone rubber [39].

Material properties of the endolymph and perilymph were assumed to be identical given their similar compositions. These properties were assigned as reported by Shen [40].

To our best knowledge, there is no published description of inner ear bone density for chinchillas. Therefore, the density of all osseous tissue was assumed to be 1200 kg/m3 as previously done by Gan in her human cochlea model with further support from Wang et al.’s conclusion that chinchilla bones have a lower density than human bones [34,41]. The Young’s modulus used for osseous tissue was 14.1 GPa as done in the human cochlea model reported by Wang et al.

### 2.5. Boundary Conditions

Simulation was carried out in Ansys. The outer bounds of the model lie in the division between the lymphatic fluids of the inner ear and the bony labyrinth. To approximate the rigidity of the bony labyrinth this outer surface was fixed. Fluid-solid interfaces were defined for each solid face in contact with endolymph or perilymph. Both the endolymph and perilymph were defined as acoustic bodies to ensure the propagation of acoustic waves. Harmonic acoustic simulation was conducted to assess the BM displacement with the displacement of the stapes footplate used as input. Experimentally determined parameters were used for the amplitude and frequency of stapes displacement at 90 dB [34]. BM displacement perpendicular to its surface was determined and normalized by stapes footplate displacement for analysis. Boundary conditions of the healthy and implanted models were identical aside from additional fluid-solid interfaces being defined on the outer surfaces of the cochlear electrode.

## 3. Results

Figure 7a shows the raw, unnormalized displacements of the basilar membrane at each tested frequency. Figure 7b shows the normalized magnitude of displacements along the cochlea. Excess noise was suppressed by applying a local filter across every 0.2 mm of the cochlea. This noise is to be expected at the overlaps of curves between two frequencies due to their differing wavelengths [43]. The majority of noise occurs towards the end of the cochlea as acoustic waves disperse, as seen in Figure 7a. This model may generate more noisy data due to the accurate triangular shape of the scala media. The magnitude of displacement of the basilar membrane decreases as frequencies become lower. This phenomenon can be explained by the heightened stiffness of the basilar membrane towards the base and has been observed in other studies [45,46,47].

**Figure 7.**
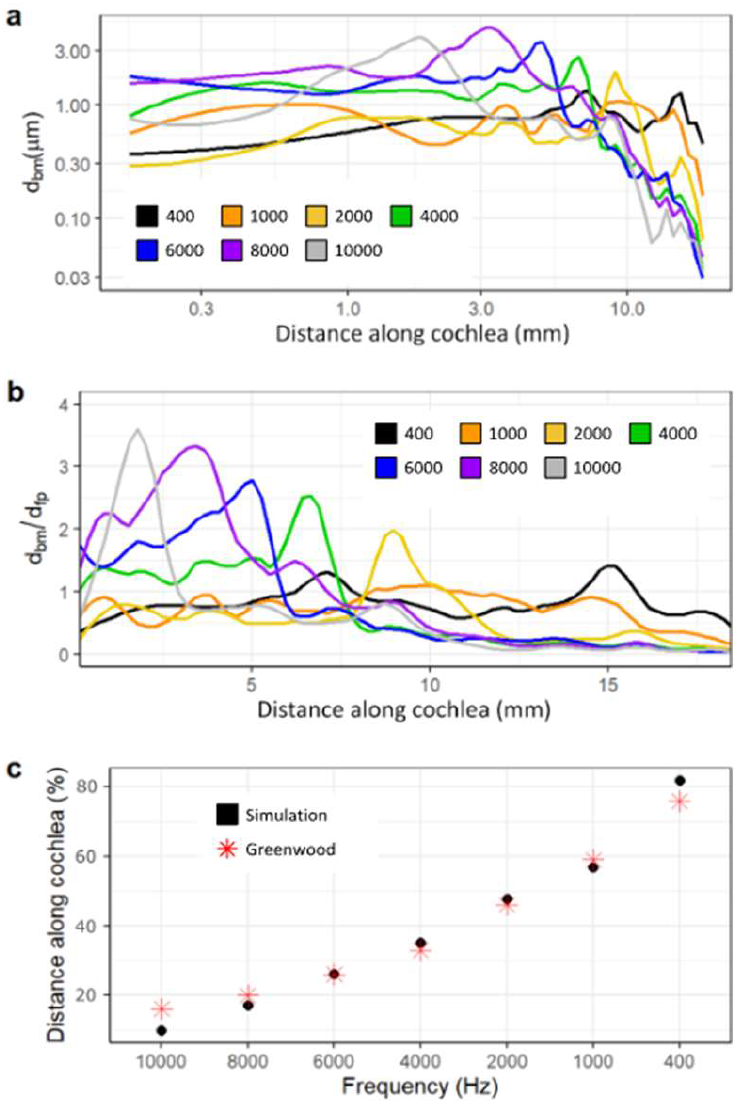
Displacement of the basilar membrane from base to apex of the cochlea before insertion of the CI. (400 Hz: black, 1000 Hz: orange, 2000 Hz: yellow, 4000 Hz: green, 6000 Hz: blue, 8000 Hz: purple, 10000 Hz: grey) Model input was experimentally determined frequency dependent displacement of the stapes at 90 dB [34]. 7a and 7b show unnormalized displacement of the basilar membrane and basilar membrane displacement normalized by that of the stapes footplate, respectively. 7c shows the location of maximum displacement for the model (black) at each frequency compared to an experimentally obtained benchmark [42].

The models’ integrity was verified by examining [42]. Figure 7c shows the tuning effect of the model, gauged by the locations of maximum displacement, in comparison to the published data [42]. This model ex the tuning effect of the cochlea and comparing it with a common source for data on the frequency and position-dependent displacement of the basilar membrane. It exhibits a mostly linear, downward trend in the magnitude of displacement as frequency decreases and is very similar in locations of maximum displacement for all assessed frequencies compared with published results [42]. This was the expected outcome, indicating that the tuning effect of the cochlea is vital to the proper perception of pitch [43, 44].

Figure 8 shows the data collected from the simulation with the cochlear implant. Results were collected for insertion angles between 180 and 900 degrees in increments of 180 degrees. The general trend in this model as the insertion angle increases is clear: CI surgery has the potential to have little effect on residual hearing. Magnitudes of displacement varied only slightly with all insertion angles and locations of maximum displacement were almost exactly consistent with the control. Only results for a 180-degree and 900-degree insertion are shown for the sake of brevity as all trials were very similar.

**Figure 8.**
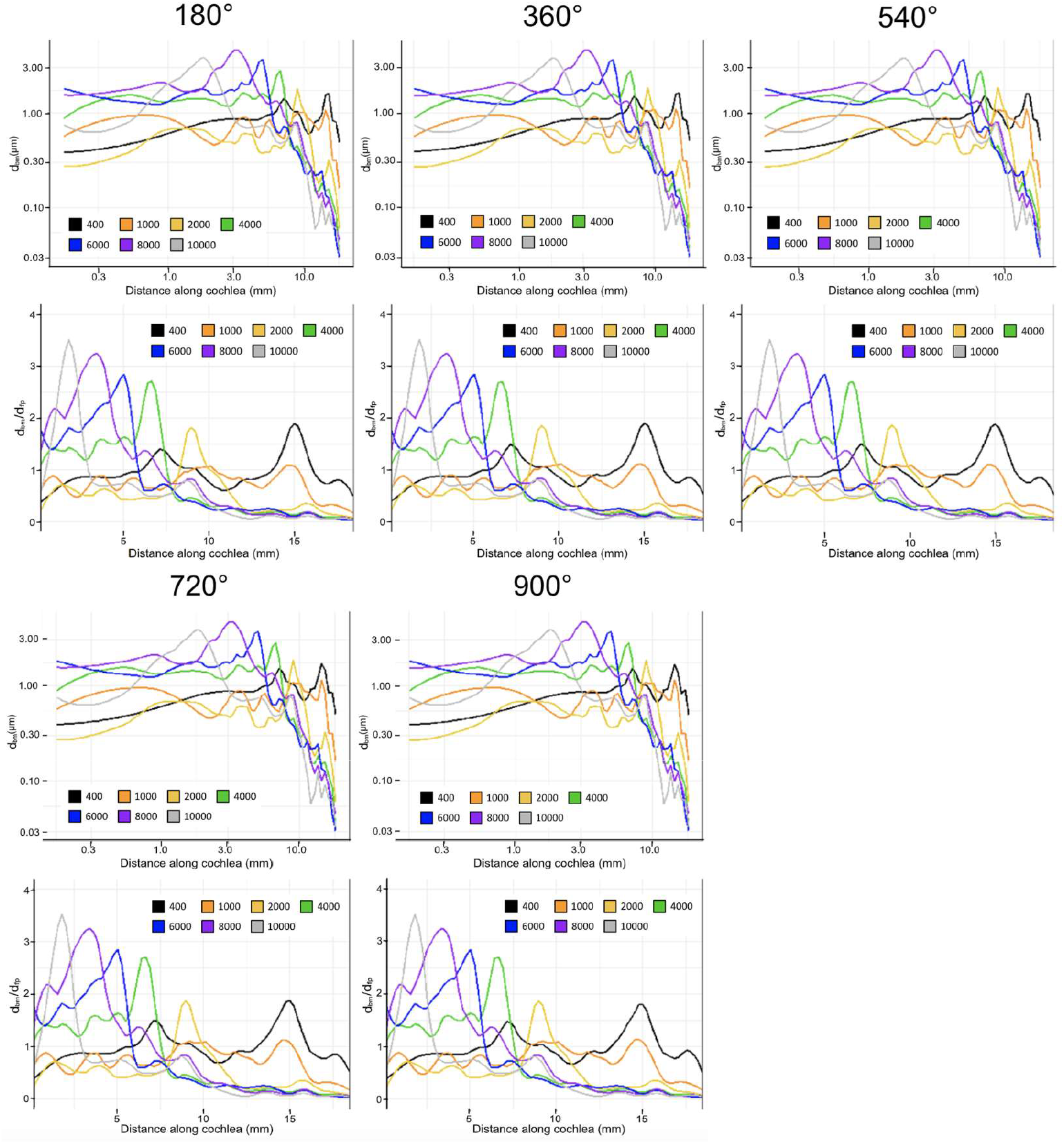
Displacement of the basilar membrane from base to apex of the cochlea at each evaluated insertion angle of the cochlear electrode. (400 Hz: black, 1000 Hz: orange, 2000 Hz: yellow, 4000 Hz: green, 6000 Hz: blue, 8000 Hz: purple, 10000 Hz: grey) Model input was experimentally determined frequency dependent displacement of the stapes at 90 dB [34]. The upper figure in each set and the lower set show unnormalized displacement of the basilar membrane and basilar membrane displacement normalized by that of the stapes footplate, respectively.

There were two major findings from these results: 1) The tuning effect of the cochlea is not significantly altered after the insertion of cochlear electrodes, representing an accurate perception of pitch. 2) The magnitudes of displacements in the basilar membrane are not significantly altered by the insertion of cochlear electrodes, representing an accurate perception of volume.

## 4. Discussion

Finite element analysis of the model in this study has two major implications: 1) Cochlear implant surgery can have minimal effect on residual hearing. 2) The insertion angle of CIs, apart from their potential to physically damage the cochlea, has little effect on residual hearing. Literature on this topic is divided. Some studies have found cochlear implant surgery to have an effect on residual hearing, like Gan’s model on the subject [34]. Others have found little to no long-term effect [45]. The general consensus is that without tip fold-over, displacement of the electrode, or some other fault, the effect of CI surgery on residual hearing is minimal [48,49]. Our results agree strongly with this conclusion. Insertion trauma is common, but it is not inevitable. While this model does represent the best-case scenario of implantation, it does not detract from the applicability to electrode design. Furthermore, as surgery techniques continue to improve and additional technology is utilized, the rate of complications is expected to decrease and thus the model will become more applicable.

It was surprising that basilar membrane displacement was almost entirely unaffected in our model regardless of the insertion angle. Iso-Mustajärvi’s 2019 study provides an explanation. He asserts that the primary contributor to the loss of residual hearing function is the stiffening of the round window membrane. Compared to simplified two-chambered or straight cochlear models, the three-chamber spiral cochlear model provides a more accurate representation of inner ear mechanics. This design introduces vital conditions which differentiate between forward and backward-driving conditions. These details also allow the model to exhibit a more realistic sensitivity to the addition of a cochlear implant than its counterparts by accurately representing distances between the electrode and surrounding structures.

It is desirable to create longer electrodes as they allow the patient to sense a wider range of frequencies more effectively [8, 46]. Results from this study support the theory that cochlear implants can have minimal effect on the mechanics of the basilar membrane, even when inserted into the apex of the cochlea. These results can inform CI electrode design; longer, more slender CI electrode designs should be prioritized to preserve residual hearing function.

There are many future applications for this model. A vestibular implant electrode could be designed and added to examine its effect on residual balance. Electrodes could be optimized with the finite element model to allow for new technologies which minimize the unintended effects of cochlear implants on surrounding nerves. This model is capable of being attached to a developed model of the chinchilla middle and outer ear. The middle ear transfer function in various scenarios could be recorded to aid clinical diagnosis of ear disorders or problems with CI function. Finally, the effect of cochlear implant surgery on residual balance could be studied using the present model and similar boundary conditions.

In the future, a series of models derived from different species will continue to be developed in our lab to enhance inner ear implantable device design and evaluation of residual hearing and balance. With the addition of the nervous system, these models will be capable of simulating electrical stimuli. Simulations in different species will assist electrode design at various stages (e.g. initial design in rodents, further optimization in primates, and clinical trials in humans).

## Author Contributions

Conceptualization, C.D.; methodology, N.C., J.L, C.M, and B.P.; software, N.C.; validation, C.D.; formal analysis, N.C.; investigation, N.C.; resources, C.D.; data curation, J.L., K.Z., and N.C.; writing–original draft preparation, N.C, C.D., C.M., B.P., and M.S.; writing–review and editing, T.E. and C.L.; visualization, T.E. and C.L.; supervision, N.C.; project administration, C.D.; funding acquisition, C.D.. All authors have read and agreed to the published version of the manuscript.

## Funding

The research was supported by University of Oklahoma Faculty Startup Fund

## Institutional Review Board Statement

A chinchillas was used for CT/MRI scanning, which was performed in accordance with a protocol approved by the University of Oklahoma Animal Care and Use Committee, which is accredited by the Association for the Assessment and Accreditation of Laboratory Animal Care (AAALAC) International and consistent with European Community Directive 86/609/EEC.

## Data Availability Statement

The data presented in this study are available on request from the corresponding author. The data are not publicly available due to policy of University of Oklahoma.

## Acknowledgements

The authors would like to acknowledge Abderrahmane Hedjoudje and Charles Della Santina at Johns Hopkins University and Rong Gan at University of Oklahoma for their generous suggestions and help for model creation and simulations.

## Conflicts of Interest

The authors declare no conflict of interest. The terms of this arrangement are being managed in accordance with University of Oklahoma policies on conflict of interest.

## Software Used

1. 3D Slicer. Version Date 4.11.2021. Open-Source. Available at https://www.slicer.org/.
2. MeshMixer. Version 3.5.474. Autodesk Inc., San Rafael, CA. Available at https://www.meshmixer.com/.
3. SpaceClaim. Version 2020.R2. Ansys Inc., Canonsburg, PA. Available at https://www.spaceclaim.com/.
4. SolidWorks. Version 2021/0128 (28-Jan-2021). Dassault Systèmes, Vélizy-Villacoublay, France. Available at https://www.solidworks.com/.
5. HyperMesh. Version Date 2017. Altair Engineering, Troy, MI. Available at https://www.altair.com/hypermesh/.
6. Ansys Workbench. Version 19.2. Ansys Inc, Canonsburg, PA. Available at https://www.ansys.com/.

## Notes

### Competing Interest Statement

The authors have declared no competing interest.

